# Proximal protein landscapes of the type I interferon signaling cascade reveal negative regulation by the E3 ubiquitin ligase PJA2

**DOI:** 10.1101/2023.10.04.560926

**Authors:** Samira Schiefer, Benjamin G. Hale

## Abstract

Deciphering the intricate dynamic events governing type I interferon (IFN) signaling is critical to unravel key regulatory mechanisms in host antiviral defense. Here, we leveraged TurboID-based proximity labeling coupled with affinity purification-mass spectrometry to comprehensively map the proximal proteomes of all seven canonical human type I IFN signaling cascade members under basal and IFN-stimulated conditions. This established a network of 103 proteins in close proximity to the core members IFNAR1, IFNAR2, JAK1, TYK2, STAT1, STAT2, and IRF9. We validated several known constitutive protein assemblies, while also revealing novel stimulus-dependent and -independent associations between key signaling molecules. Functional screening further identified PJA2 as a negative regulator of IFN signaling via its E3 ubiquitin ligase activity. Mechanistically, PJA2 interacts with the Janus kinases, TYK2 and JAK1, and promotes their non-lysine and non-degradative ubiquitination, resulting in restrained downstream STAT signaling. Our high-resolution proximal protein landscapes provide global insights into the type I IFN signaling network, and serve as a valuable resource for future exploration of its functional complexities.

## Introduction

Virus-recognition by infected cells triggers the production of multiple cytokines, including the antiviral type I interferons (IFNs). These type I IFNs (e.g. IFN-α or IFN-β) act by upregulating the expression of antiviral genes (IFN-Stimulated Genes; ISGs) to restrict virus replication, but they also act to balance the innate immune system and to promote the adaptive immune response^1,2^. IFNα/β receptor 1 (IFNAR1) and IFNAR2 form a heterodimeric complex on the surface of cells to recognize secreted type I IFNs and to signal via the classical Janus kinase / Signal Transducer and Activator of Transcription (JAK/STAT) pathway to induce antiviral effects. More precisely, high affinity binding of type I IFNs to IFNAR2 promotes the dimerization of both receptor chains and permits activation of the receptor-associated intracellular kinases Janus kinase 1 (JAK1) and Tyrosine kinase 2 (TYK2)^3,4^. These kinases trans-/auto-phosphorylate, and in turn phosphorylate STAT1 and STAT2 on specific tyrosine residues, leading to STAT1/2 translocation into the nucleus where they assemble with Interferon Regulatory Factor 9 (IRF9) to form the IFN-Stimulated Gene Factor 3 (ISGF3) complex. This transcription factor complex then binds to IFN-Stimulated Response Elements (ISREs) in ISG promoter regions and, with the coordinated action of chromatin-remodeling complexes, activates the expression of many hundreds of ISGs^5–7^. The protein products of ISGs play key roles in limiting virus spread by various potent antiviral mechanisms^2,8^.

Dysregulation of the type I IFN signaling cascade is associated with a variety of pathological conditions in humans. For instance, autoimmune diseases (such as systemic lupus erythematosus) can be caused by IFN system overactivation^9–11^, while increased susceptibility to severe viral diseases or chronic viral infections can be caused by IFN defects^12^. Thus, type I IFN signaling has to be tightly and intricately regulated with multiple redundant mechanisms, which together help to limit potential overactivation while still being able to react rapidly and effectively against invading viral pathogens^13^. While we do not have a full understanding of the complexities of type I IFN signaling network regulation, it is clear that protein post-translational modifications, such as (de-)phosphorylation, ubiquitination, SUMOylation, and acetylation can be key players^14^. For example, IFNAR1 is phosphorylated by the IFN-inducible Protein Kinase D2 (PKD2) on a degron motif leading to its ubiquitination, internalization and lysosomal degradation in order to prevent hyperstimulation^15–17^. In addition, phosphatases, including T Cell Protein Tyrosine phosphatase (TC-PTP) and Protein-Tyrosine Phosphatase 1B (PTP1B), have been reported to dephosphorylate and inhibit JAK1 and TYK2^18,19^. Furthermore, the Suppressors Of Cytokine Signaling 1/3 (SOCS1/3) are negative regulatory proteins that inhibit JAK1/TYK2^20,21^ by sterically hindering their kinase activities by blocking substrate binding^22^, but have also been described to induce their ubiquitin-mediated degradation^23^. Similarly, Ubiquitin Specific Peptidase 18 (USP18) interacts with IFNAR2/STAT2 and reduces JAK1 phosphorylation and activity^24^. At a different level, the Protein Inhibitor of Activated STAT (PIAS) proteins use diverse mechanisms to inhibit IFN signaling^25^. A notable example is PIAS1, an E3-type Small Ubiquitin-like Modifier (SUMO) ligase, that interferes with STAT1 by SUMOylating it and preventing STAT1 from binding to the promoter region of specific ISGs^26–28^. Finally, STAT2 and IRF9 can be acetylated by CREB-binding protein (CBP) in their DNA-binding domains in response to IFN stimulation to ensure proper ISGF3 activity^29^. The identification and molecular characterization of cellular factors that regulate type I IFN signaling has therefore proven to be invaluable in unraveling the functionality of this system. Such work has also previously provided key insights into potential therapeutic targets to treat diseases associated with dysregulation of the IFN system, with key examples being approved drugs such as JAK inhibitors^30–32^.

Our current knowledge on type I IFN signaling regulation mainly comes from factor-specific protein interaction studies (e.g. yeast two-hybrid^33–35^ screens and immunoprecipitations^36^), or from genetic studies using chemical mutagenesis^37^, siRNAs^38^ or CRISPR/Cas9-based^39,5^ methods. However, conventional biochemical interaction studies can suffer from biases towards the enrichment of very strong interactors, and may miss weak, transient, or even cell context-dependent factors. Gene depletion studies also favor the identification of genes non-essential for cell viability at the expense of equally important factors that may have additional critical cellular roles. In this regard, recent advances in proximity labeling technologies have revealed their potential as powerful tools for deciphering the complex protein environments that underpin cellular signaling networks^40,41^. A major benefit is the possibility to uncover cell context-dependent low-affinity interactions and stimulus-dependent transient interactions in minute-scale temporal resolution^42^. In this study, we therefore aimed to leverage TurboID-based proximity labeling^43^, coupled with affinity purification mass spectrometry (AP-MS), to produce an unprecedented global glimpse into the dynamic proximal proteomes associated with the seven core members of the human type I IFN signaling cascade: IFNAR1, IFNAR2, JAK1, TYK2, STAT1, STAT2, and IRF9. By attempting to capture temporal changes in protein interactions following type I IFN stimulation, we sought to unveil novel regulatory mechanisms and transient associations that help to orchestrate type I IFN signaling precision. Here, we present high-resolution landscapes that include 103 proteins closely associated with the core type I IFN signaling machinery, revealing both known protein assemblies and novel unappreciated complexes. By integrating results from a functional screening approach, we focused on dissecting the role of PJA2 as a previously unrecognized negative regulator of IFN signaling through its E3 ubiquitin ligase activity. PJA2 could promote the non-lysine and non-degradative ubiquitination of Janus kinases to limit downstream STAT signaling. Thus, our study offers a comprehensive exploration of the individual proximal proteomes associated with the type I IFN signaling cascade and provides an enriched framework to expand our current understanding of this pathway.

## Results

### System-wide identification of the type I IFN signaling proximal proteome

Rapid promiscuous biotinylation of proximal (∼ 35 nm) proteins within minutes makes TurboID an ideal enzyme to study cell signaling events involving low affinity or temporally regulated transient interactions^43,44^. We therefore sought to apply TurboID-based proximity labeling and AP-MS analysis to identify proteins in proximity to each of the seven canonical human type I IFN signaling cascade components during IFN-α2 stimulation. Using lentiviral vectors, we engineered eleven independent hTERT-immortalized human lung fibroblast (MRC-5) cell lines to stably express TurboID-tagged IFNAR1, IFNAR2, JAK1, TYK2, STAT1, STAT2 or IRF9, as well as TurboID-tagged localization controls based on GFP alone or fused to the Lyn11 plasma membrane localization signal^45^ or classical nuclear export^46^ or nuclear localization^47^ signal sequences (**Fig. 1a, Supplementary Fig. 1a**). Immunofluorescence microscopy confirmed the homogenous expression and correct intracellular localization of each TurboID-tagged protein (**Supplementary Fig. 1b**). Furthermore, endogenous IFN signaling component depletion revealed the functionalities of each TurboID-tagged IFN signaling member, with all TurboID-tagged components retaining appreciable activity (**Supplementary Fig. 1c**). Kinetic analysis of IFN-α2 signaling in MRC-5 cells revealed that critical events such as STAT1/STAT2 phosphorylation and STAT1 nuclear translocation all occurred between 10 min and 120 min post IFN-α2 stimulation (**Supplementary Fig. 1d, e**), suggesting that timepoints within this window are suitable to identify functionally relevant proteins in proximity to each IFN signaling component. We consequently stimulated each of our engineered cell lines with IFN-α2 for various times (20 – 120 min) and supplemented the culture medium with exogenous biotin 15 min prior to harvesting in order to biotinylate proximal proteins. After affinity purification of the biotinylated proteins, enriched factors were detected and quantified by label-free mass spectrometry (**Fig. 1a**). Proteins with high probability of being proximal to each IFN signaling component at each timepoint post IFN-α2 stimulation were identified by comparing label-free protein quantities against the similarly localized GFP control across three independent replicates with Significance Analysis of Interactome (SAINT)^48,49^. Using stringent selection criteria (4-fold enrichment over the similarly localized GFP control, 2-fold enrichment over all other GFP controls, and an interaction probability of at least 70%), our analyses yielded a total of 103 proteins in proximity to IFNAR1/2, JAK1, TYK2, STAT1/2, and IRF9, which does not include the seven canonical IFN signaling components themselves that were also identified (**Fig. 1b, c, Supplementary Data 1**). Consistent with previous literature, our data confirmed the constitutive interaction between STAT1 and STAT2^50^, and the establishment of a specific STAT2-IRF9 complex lacking STAT1 in unstimulated cells^51^ (**Fig. 1d**). Additionally, our method highlighted the transient IFN-α2 stimulated interaction between TYK2 and STAT1/STAT2, as well as between STAT1 and IRF9^52,53^ (**Fig. 1d**). Further supporting the overall validity of our newly-identified type I IFN signaling proximal proteome, several of the proximal proteins could be confirmed by immunoblot analysis of independent small-scale proximity labeling experiments: e.g. the proximity of IL6ST with JAK1 and TYK2 (**Fig. 1e**), DNAJA2 with STAT1 and STAT2 (**Fig. 1f**), CNOT1 with IRF9 (**Fig. 1g**), and the IFN-α2-stimulated proximity of Kinesin Light Chain 2 (KLC2) with STAT2 (**Fig. 1h**). Thus, our TurboID-based study has generated a robust dataset of >100 proteins with high probability of being in proximity to the seven canonical IFN signaling cascade components. This dataset will be an essential resource for future exploration of novel functional aspects of this system.

**Fig. 1:**
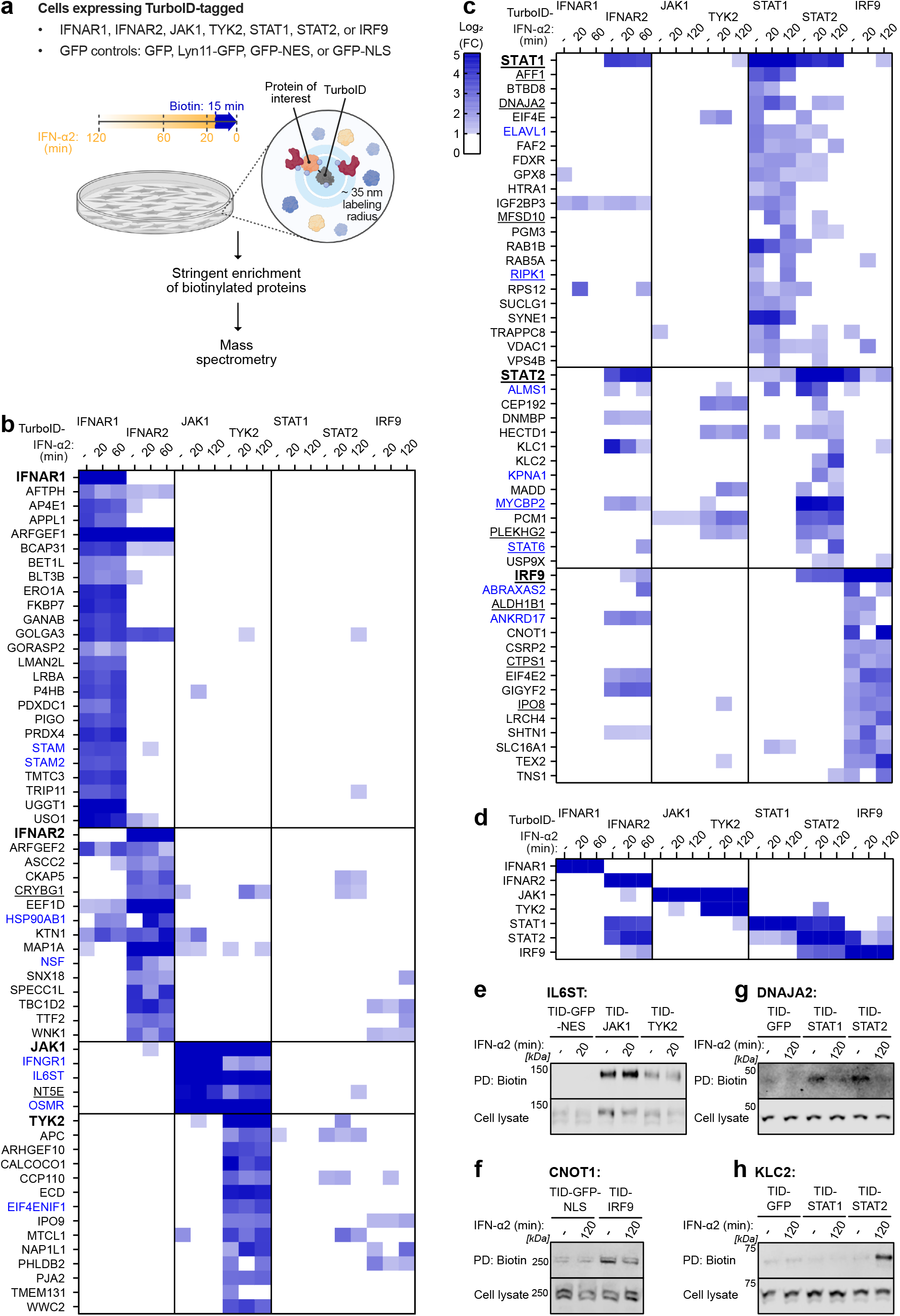
System-wide identification of the type I IFN signaling proximal proteome. **a** Schematic representation of the type I IFN signaling TurboID screening strategy. The indicated MRC-5/hTERT cell lines were stimulated with 1000 IU/mL IFN-α2 for 120, 60, 20, or 0 min, and 500 μM biotin was added for the final 15 min prior to harvesting. Biotinylated proteins were purified with streptavidin beads prior to identification and quantification by mass spectrometry. Three biologically independent replicates were performed. **b** - **d** Heat map representations of log_2_(fold change, FC) in abundance of proteins identified for each of the indicated TurboID constructs at each timepoint post IFN-α2 stimulation. Abundance was determined by label-free quantification and made relative to the relevant background control (i.e. the similarly localized TurboID-GFP construct). Shown are all proteins that were considered significant according to our selection criteria and that were enriched >4-fold in at least one condition (see Methods). Only log_2_(FC) > 1 are depicted. **b**, **c** Data for all identified proteins. **d** Data focused only on canonical type I IFN signaling members. In **b**, **c**, previously annotated interactors of canonical type I IFN signaling components according to BioGRID^54^ and/or STRING^55^ are labeled in blue, while previously identified ISGs^58,59^ (>1.5-fold with IFN) are underlined. **e** - **h** Immunoblot confirmation of four newly identified type I IFN signaling proximal proteins from independent small-scale TurboID experiments performed as described in **a**. Total cell lysates and enriched biotinylated proteins (pull-down, PD) from the indicated conditions were immunoblotted for IL6ST (**e**), CNOT1 (**f**), DNAJA2 (**g**) and KLC2 (**h**).

### Global analysis of the type I IFN signaling proximal proteome

Bioinformatic analysis of our type I IFN signaling proximal proteome revealed an interconnected network of factors linked both physically and functionally (**Fig. 2**). Striking proportions of the factors we identified have previously been associated with components of the type I IFN signaling response (16%, 16 of 103; according to BioGRID^54^ and/or STRING^55^). In addition, 13% (13 of 103) are known to be involved in viral processes^56,57^, and 12% (12 of 103) are annotated as ISGs^58,59^. Notably, 83% of these ISGs identified in our study are in close proximity to the transcription factor components, STAT1, STAT2 and IRF9. Among the 16 proximal proteins that have been previously linked to type I IFN signaling were STAM and STAM2, which we identified in proximity to IFNAR1 and that have very recently been shown to inhibit IFNAR1-associated TYK2 function in unstimulated cells^60^. In addition, ELAVL1, which we found in proximity to STAT1, is an RNA-binding protein that stabilizes the mRNAs of ISGs^61^. Furthermore, ABRAXAS2, which we identified proximal to IRF9, is a subunit of the BRCC36 isopeptidase complex (BRISC), and is important for ensuring proper STAT1 phosphorylation in response to herpes simplex virus infection^62^. Another factor we identified proximal to IRF9, GIGYF2, appears to be a repressor of ISG activation in the absence of IFN^39^. Additional gene ontology^56,57^ analysis of the proteins we identified in proximity to type I IFN signaling components using DAVID^63^ revealed significant enrichment in factors associated with plausibly relevant biological processes for IFN signaling, such as ‘protein transport’, ‘cytokine-mediated signaling’ and ‘protein import into nucleus’ (**Supplementary Fig. 2a-c, Supplementary Data 2**). In this context, it is tempting to speculate on the possible trafficking role of factors such as KLC2, which we could clearly validate as a protein in close proximity to STAT2 after IFN-α2 stimulation (**Fig. 1h**), given its known function in binding specific cargoes that have stimulus-dependent cytosol to nucleus signaling activity via microtubules^64^.

**Fig. 2:**
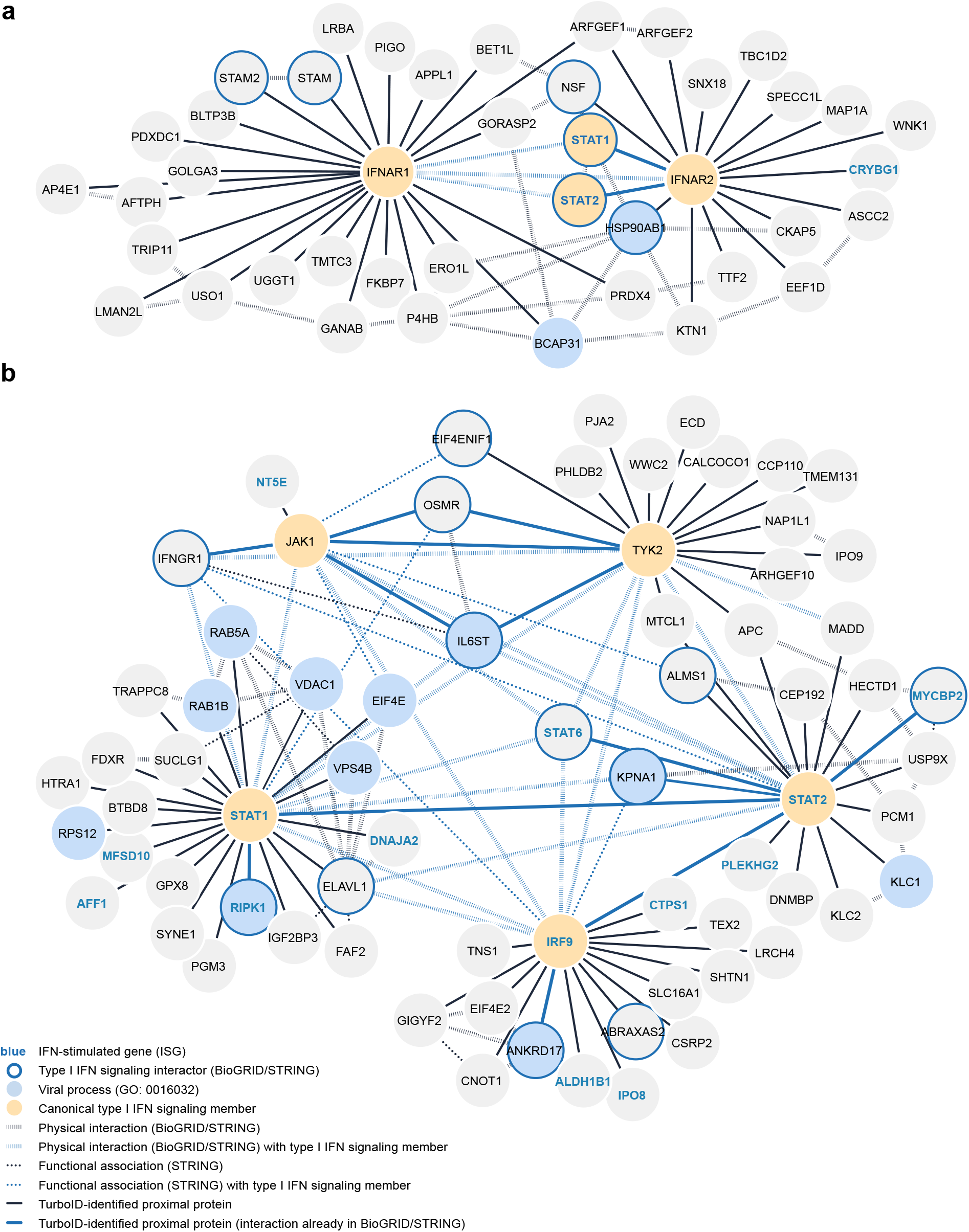
Global analysis of the type I IFN signaling proximal proteome. **a** Visualization of the interaction network of proteins identified in proximity to IFNAR1 and IFNAR2. Previously annotated interactions are depicted. **b** Visualization of the interaction network of proteins identified in proximity to JAK1, TYK2, STAT1, STAT2, and IRF9. Previously annotated interactions that were not found in our TurboID screen are depicted where they link proteins found in proximity to the same type I IFN signaling component, or towards JAK1, TYK2, STAT1, STAT2, or IRF9. For **a** and **b**, physical interactions and functional associations annotated on BioGRID^54^ and/or STRING^55^, known ISGs, factors involved in viral processes, and the proximal proteins identified in this study are all highlighted as indicated in the figure key. Networks were generated using Cytoscape 3.9^101^.

### siRNA screening identifies proximal proteins that functionally regulate type I IFN signaling

To prioritize the newly identified proximal proteins that influence type I IFN signaling, we used siRNA pools to deplete MRC-5 cells of 50 high confidence candidates one by one, and subsequently assessed IFN-α2-stimulated (4 h and 16 h) antiviral activity against the IFN-sensitive VSV-GFP reporter virus (**Fig. 3a**). As controls, we depleted IFNAR2 and IRF9 (two positive regulators of type I IFN signaling), as well as USP18 and SOCS3 (two negative regulators of type I IFN signaling). We initially classified factors as important for type I IFN signaling if their depletion increased or decreased IFN-α2-stimulated antiviral activity by at least 2-fold compared to the non-targeting control siRNA (NTC) in two independent screening runs. Using these criteria, we could confirm IFNAR2 and IRF9 as positive regulators of type I IFN signaling, and SOCS3 and USP18 as negative regulators of type I IFN signaling (**Fig. 3b, c, d**). In addition, depletion of 16 out of the 50 candidates reproducibly increased the sensitivity of VSV-GFP to 4 h and/or 16 h IFN-α2 stimulation, similar to USP18 and SOCS3 depletion, suggesting that they may also act as negative regulators of type I IFN signaling (**Fig. 3d, Supplementary Fig. 3, Supplementary Data 3**). These regulators included USP9X, which we identified in proximity to STAT2 in our proteomics screen, and that has not previously been associated with type I IFN signaling, but has been reported to play a role in the closely related JAK2-dependent JAK/STAT signaling pathway^65^. To assess whether any of the newly identified candidate regulators altered the ability of IFN-α2 to stimulate antiviral gene expression directly, we again used siRNA pools to deplete a subset of the candidates in MRC-5 cells, and assessed IFN-α2-stimulated *ISG54* and *MX1* mRNA expression levels at 4 and 16 h respectively, which were the peak timepoints for individual expression of these two ISGs. As expected, depletion of IFNAR2 or IRF9 led to significant decreases in IFN-α2-stimulated expression of these ISGs, while depletion of USP18 and SOCS3 led to increases in their expression (**Fig. 3e, f**). Out of the candidates tested in this assay, depletion of three (CNOT1, PJA2, and IGF2BP3) led to a significant and robust increase in IFN-α2-stimulated expression of *ISG54* or *MX1* mRNAs (**Fig. 3e, f**), further confirming their role as negative regulators with likely direct effects on type I IFN signaling. Other factors, such as AFF1, ALMS1, NT5E, OSMR, TBC1D2 and USP9X showed effects as potential negative regulators of type I IFN signaling with a 50% increase in *ISG54* or *MX1* mRNAs after depletion of these factors. However, these effects did not reach statistical significance in the assays we employed. Interestingly, of the significant factors, CNOT1 (which we identified in proximity to IRF9) is part of the CCR4-NOT complex that has previously been reported to negatively regulate type I IFN signaling^66,67^. These results indicate that several members of the type I IFN signaling proximal proteome play functionally relevant roles in this pathway.

**Fig. 3:**
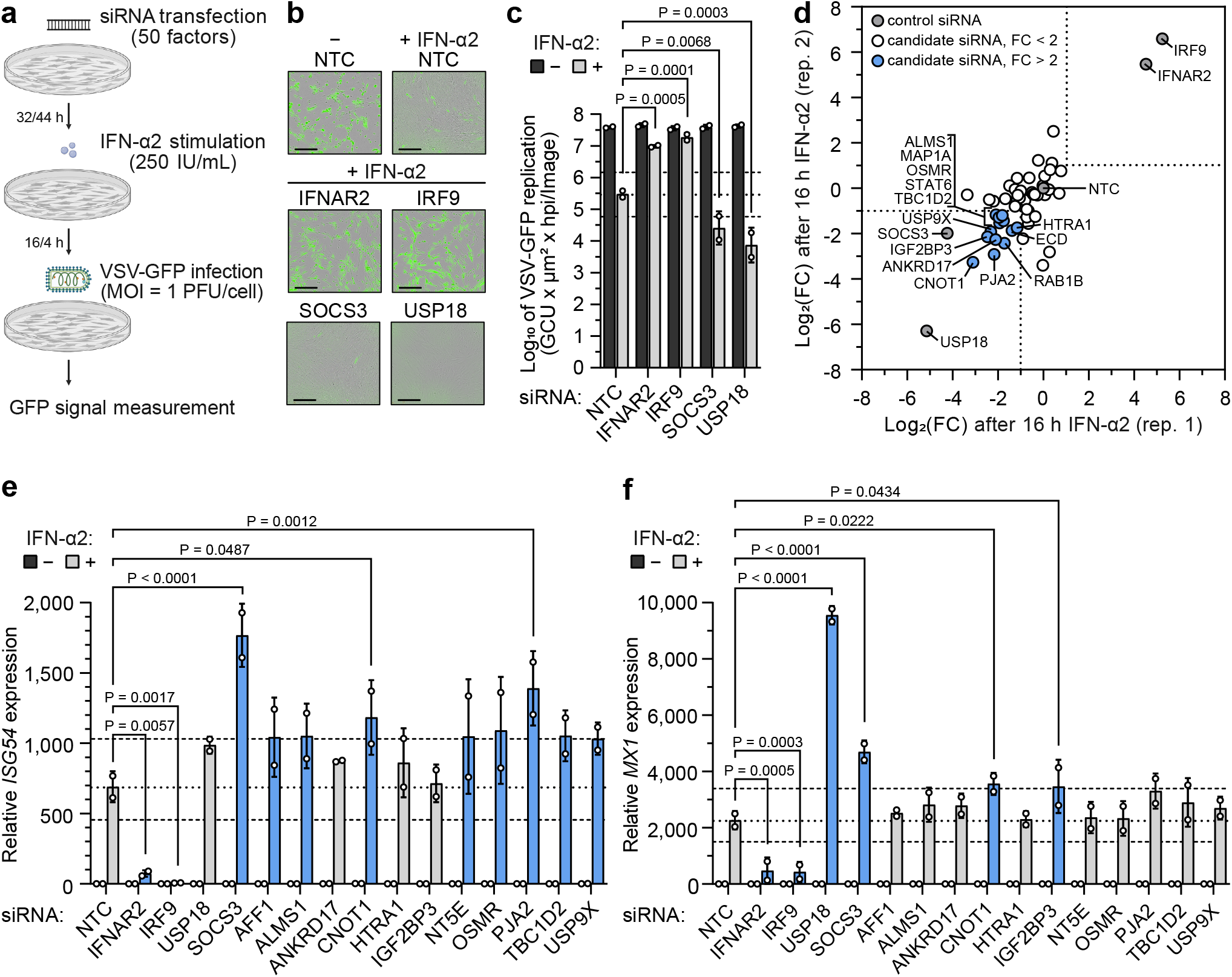
siRNA screening identifies proximal proteins that functionally regulate type I IFN signaling. **a** Schematic representation of the VSV-GFP based siRNA screening approach. siRNA pools targeting 50 factors were individually reverse transfected into MRC-5/hTERT cells prior to stimulation with 250 IU/mL IFN-α2 for 4 or 16 h. Cells were then infected with VSV-GFP (MOI = 1 PFU/cell) and GFP signal was monitored with the IncuCyte live-cell imaging system as a surrogate readout for viral replication. **b**, **c** VSV-GFP assay controls showing viral replication in the indicated siRNA-transfected conditions ± 16 h IFN-α2 stimulation. **b** consists of representative GFP images of infected cells at 48 h post infection. Scale bars represent 300 μm. **c** shows total GFP levels calculated from area under the curve (AUC) values for VSV-GFP replication during the course of the experiment. Bars represent mean values from two biologically independent experiments. Error bars represent Standard Deviations (SDs). The dotted line indicates VSV-GFP replication in the non-targeting siRNA control (NTC) condition after IFN-α2 stimulation, and the dashed line indicates a 5-fold change. Statistically significant P values are shown and were determined by two-way ANOVA and Šídák’s multiple comparisons. **d** Log_2_(FC) in VSV-GFP replication (AUC values) after 16 h IFN-α2 stimulation of cells transfected with siRNAs targeting the indicated genes. Log_2_(FC) is relative to VSV-GFP replication in the non-targeting siRNA control (NTC) condition. Two biologically independent replicates are plotted. The dotted lines indicate a 2-fold change in VSV-GFP replication as compared to the NTC condition. **e**, **f** *ISG54* (**e**) or *MX1* (**f**) mRNA expression as determined by RT-qPCR following ± 4 h (**e**) or ± 16 h (**f**) IFN-α2 stimulation (250 IU/mL). Like the experiment described in **a**, MRC-5/hTERT cells had previously been reverse transfected with siRNA pools. Data are normalized to GAPDH expression levels in the same sample and made relative to the NTC minus IFN-α2 condition. Bars represent means ± SDs from two biologically independent experiments conducted in technical duplicates. Statistically significant P values are shown and were determined by two-way ANOVA and Šídák’s multiple comparisons. Non-significant values (P > 0.05) are not shown. The dotted line indicates ISG expression in the NTC condition after IFN-α2 stimulation, and the dashed line indicates a 1.5-fold change from the NTC. Bars with a > 1.5-fold change are colored in blue.

### PJA2 interacts with TYK2 and JAK1 and negatively regulates type I IFN signaling in an E3 ubiquitin ligase activity-dependent manner

We focused follow-up studies on PJA2, a RING-type (Really Interesting New Gene) E3 ubiquitin ligase that was first identified for its ability to control Protein Kinase A (PKA) stability by ubiquitination^68^. PJA2 has since been described to be involved in other signaling pathways such as the Hippo cascade or Wnt/β-catenin signaling, and has been reported to act either by inducing target degradation or by adding atypical ubiquitin chain linkages to regulate target function^69–71^. PJA2 currently has no known associations with IFN signaling, but we identified it in proximity to TYK2 (**Fig. 1b, Supplementary Data 1**), and our siRNA-mediated depletion studies identified it as a potential negative regulator of IFN-α2-stimulated gene expression and antiviral activity (**Fig. 3d, e, and Supplementary Fig. 3**). We confirmed that these functional effects were not limited to MRC-5 cells by performing similar PJA2 depletion studies in human A549 cells, and found that PJA2 depletion (similar to depletion of SOCS3) enhanced IFN-α2-stimulated antiviral activity in this cell line (**Fig. 4a**). In addition, we investigated whether the identified proximity of PJA2 with TYK2 could be due to a physical interaction by performing co-immunoprecipitation experiments with tagged forms of TYK2 and PJA2. A V5-tagged TYK2 construct could specifically precipitate endogenous PJA2 (**Fig. 4b**), while a Flag-tagged PJA2 construct could specifically precipitate endogenous TYK2 (**Fig. 4c**). These data reveal that TYK2 and PJA2 can physically associate with one another in the same complex, suggesting a relatively stable interaction had been identified in the TurboID-based screen. Interestingly, we found that only the related kinase, JAK1, was additionally able to co-precipitate PJA2 when all seven tagged type I IFN signaling components were tested for their interaction with the E3 ubiquitin ligase (**Fig. 4d**), revealing specificity of PJA2 interaction with the Janus kinases TYK2 and JAK1.

**Fig. 4:**
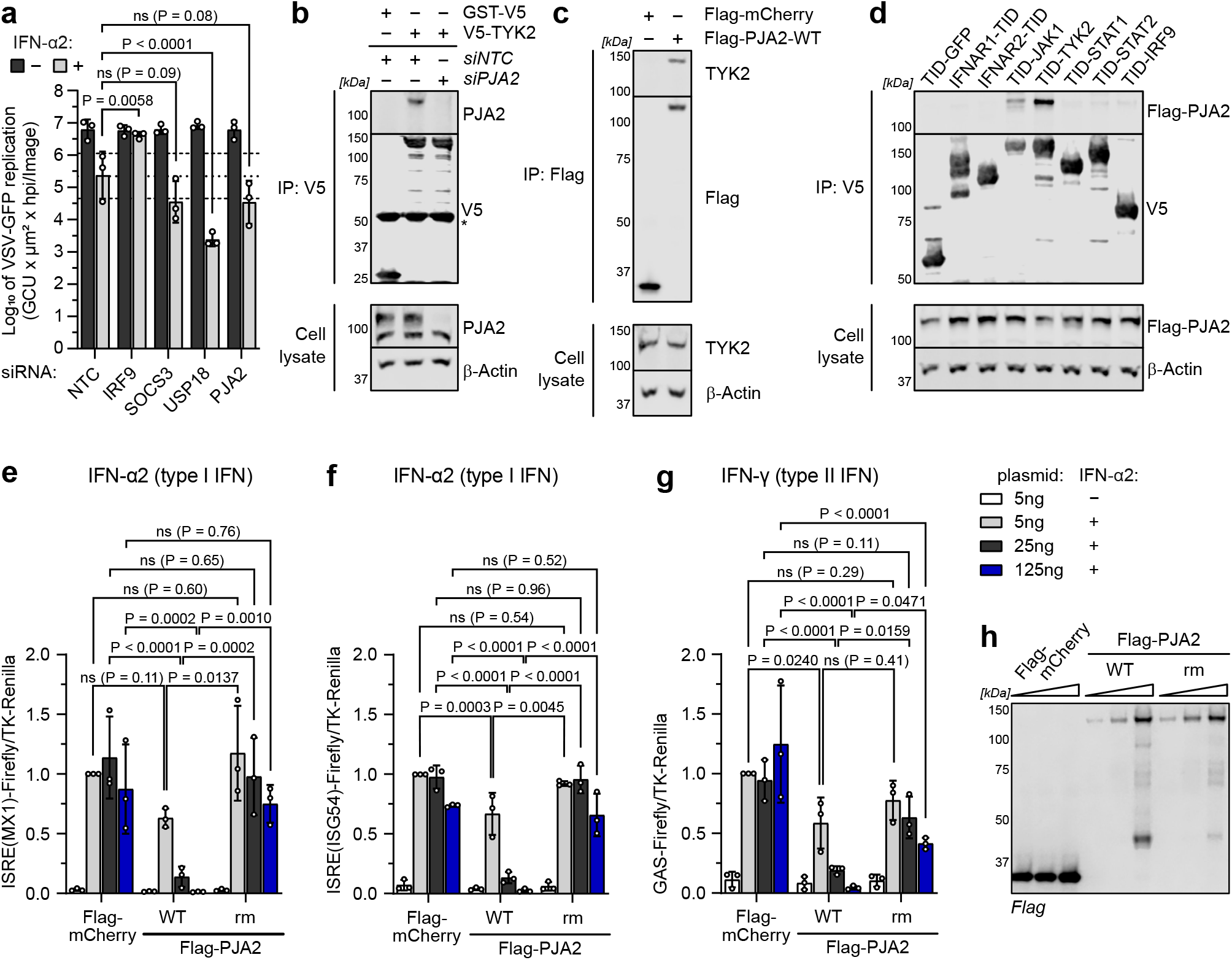
PJA2 interacts with TYK2 and JAK1 and negatively regulates type I IFN signaling in an E3 ubiquitin ligase activity-dependent manner. **a** VSV-GFP assay showing viral replication in the indicated siRNA-transfected A549s ± 16 h IFN-α2 (250 IU/mL) stimulation. Total GFP levels were calculated from AUC values for VSV-GFP replication during the course of the experiment. Means ± SDs from three biologically independent experiments are shown. The dotted line indicates VSV-GFP replication in the NTC condition after IFN-α2 stimulation, and the dashed line indicates a 5-fold change. P values were determined by two-way ANOVA and Šídák’s multiple comparisons. Non-significant (ns) values are P > 0.05. **b** HEK293T cells were co-transfected with plasmids expressing V5-TYK2 or GST-V5 together with NTC or PJA2-targeting siRNAs prior to generation of cell lysates and anti-V5 immunoprecipitation (IP). Cell lysate and IP fractions were analyzed by SDS-PAGE and immunoblotting for the indicated proteins. Data are representative of at least two biologically independent experiments. * indicates the IgG heavy chain. **c** HEK293T cells were transfected with plasmids expressing Flag-mCherry or Flag-PJA2 prior to generation of cell lysates and anti-Flag IP. Cell lysate and IP fractions were analyzed by SDS-PAGE and immunoblotting for the indicated proteins. Data are representative of three biologically independent experiments. **d** HEK293T cells were co-transfected with plasmids expressing Flag-tagged PJA2 and each of the indicated TurboID-V5-tagged type I IFN signaling components (or GFP) prior to generation of cell lysates and anti-V5 IP. Cell lysate and IP fractions were analyzed by SDS-PAGE and immunoblotting for the indicated proteins. Data are representative of three biologically independent experiments. **e** – **g** HEK293T cells were co-transfected with ISRE(MX1)-Firefly (**e**), ISRE(ISG54)-Firefly (**f**) or GAS-Firefly (**g**) plasmids together with a TK-Renilla control and 5, 25, or 125 ng of plasmid expressing Flag-tagged PJA2-WT, PJA2-rm or mCherry. Following 16 h of 1000 IU/mL IFN-α2 (**e**), 100 IU/mL IFN-α2 (**f**), or 1000 IU/mL IFN-γ (**g**) stimulation, luciferase activities were determined and normalized to the 5 ng mCherry control condition. Bars represent mean ± SD values from three biologically independent experiments. P values were determined by two-way ANOVA and Tukey’s multiple comparisons. Non-significant (ns) values are P > 0.05. **h** Representative immunoblot from the experiments shown in **e** – **g** to show the expression levels of the indicated Flag-tagged proteins. Data are representative of at least three biologically independent experiments.

We next sought to determine whether the E3 ubiquitin ligase activity of PJA2 contributes to its negative regulation of type I IFN signaling. In an orthogonal approach to our siRNA-based depletion studies, we found that overexpression of wild type (WT) Flag-tagged PJA2 led to a dose-dependent reduction in IFN-α2-induced ISRE activity in two separate (MX1 and ISG54) luciferase-based reporter assays (**Fig. 4e, f**). Interestingly, overexpression of a previously described PJA2 RING domain mutant (C634A, C671A that lacks E3 ubiquitin ligase activity^68^; PJA2-rm) was unable to reduce IFN-α2-induced ISRE activity (**Fig. 4e, f**), revealing the importance of PJA2 E3 ubiquitin ligase activity for the negative regulation of type I IFN signaling. Given that JAK1 plays an essential role in both type I and type II IFN signaling, while TYK2 activity is restricted to type I IFN signaling^4^, we also assessed the impact of PJA2 overexpression on IFN-γ-induced gamma-activated site (GAS) luciferase reporter activity. Consistent with the observation that PJA2 can interact with JAK1, overexpression of PJA2-WT led to a dose-dependent reduction in IFN-γ-induced GAS activity (**Fig. 4g**). Surprisingly, the catalytically inactive PJA2-rm also displayed a dose-dependent reduction of GAS activity, albeit to a weaker extent than PJA2-WT (**Fig. 4g**). Immunoblot analysis confirmed that PJA2-WT and the catalytically inactive PJA2-rm expressed to similar levels in these assays (**Fig. 4h**), suggesting that differences in abundance do not account for observed functional effects. These data reveal the ability of PJA2 to physically interact with the Janus kinases, JAK1 and TYK2, and identify a negative regulatory role for PJA2 during type I and type II IFN signaling processes. The ability of PJA2 to inhibit type I IFN signaling is strongly dependent on its E3 ubiquitin ligase function.

### PJA2 promotes ubiquitination of TYK2 in a lysine-independent manner

To further understand the molecular mechanism by which PJA2 negatively regulates IFN signaling, we investigated whether PJA2 could promote ubiquitination of JAK1 and/or TYK2. In cell-based ubiquitination assays, we found that V5-tagged JAK1 and TYK2 were more strongly ubiquitinated in the presence of PJA2-WT than in the presence of PJA2-rm, while a similarly tagged control protein (GST-V5) was not ubiquitinated (**Fig. 5a-c**). We also verified that untagged TYK2 is more strongly ubiquitinated in the presence of PJA2-WT than in the presence of PJA2-rm to exclude any artefactual effects arising from the use of an N-terminal V5 tag on the Janus kinases (**Fig. 5d**). These data suggest that PJA2 can indeed promote ubiquitination of JAK1 and TYK2. In support of a possible direct effect, we found that both PJA2-WT and PJA2-rm maintain their interactions with TYK2 and JAK1 (**Fig. 5e, Supplementary Fig. 4a**), indicating that loss of target ubiquitination by PJA2-rm is not due to impaired target recruitment. Furthermore, using a variety of truncation mutants, we determined that the RING domain of PJA2 is not required for its interaction with TYK2, and instead PJA2 requires a region from amino acids 531 to 630 for a strong interaction with TYK2 (**Fig. 5e, f**). This region was previously published to be required for its interaction with the regulatory subunit of another kinase, PKA^68^.

**Fig. 5:**
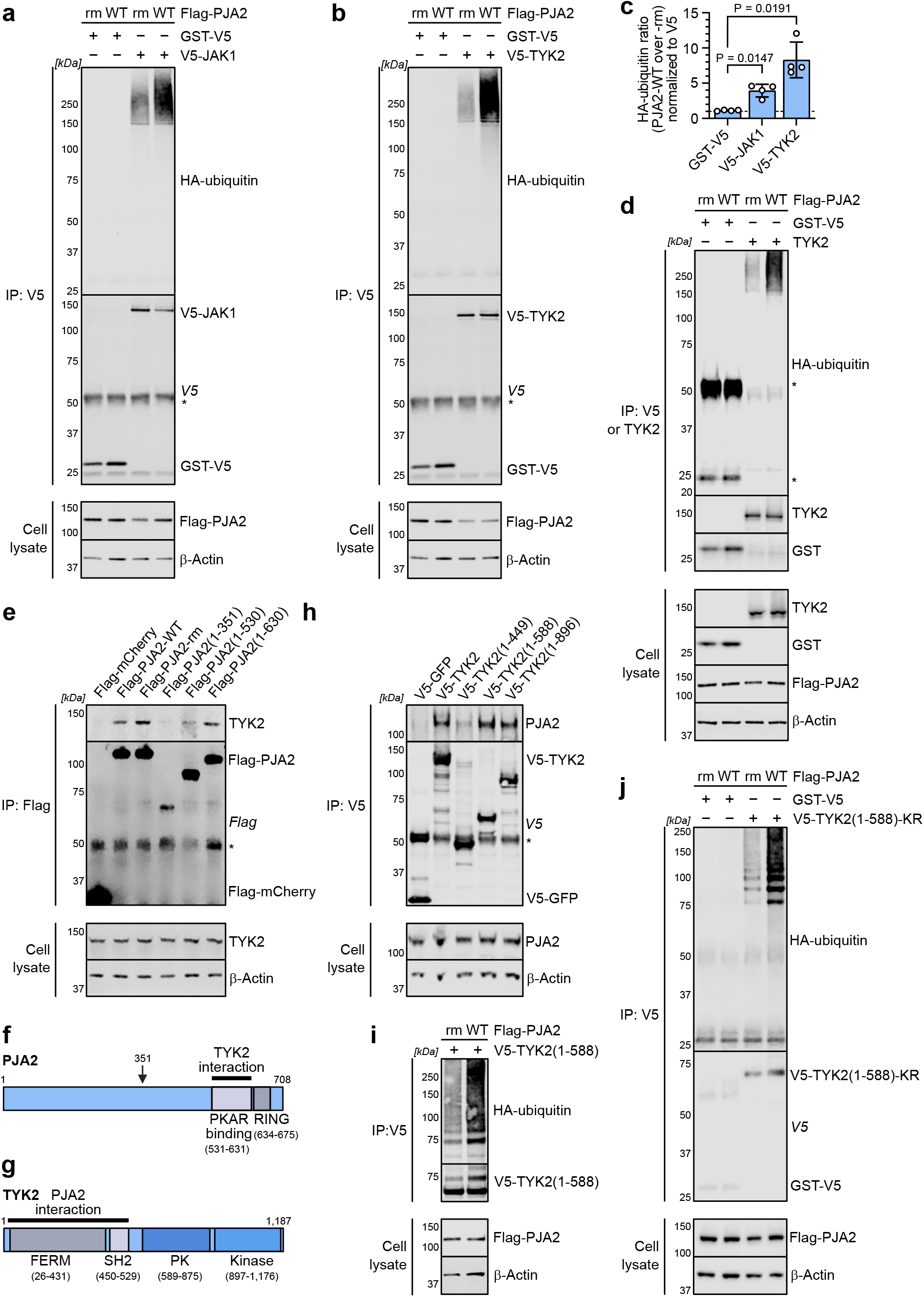
PJA2 promotes ubiquitination of TYK2 in a lysine-independent manner. **a**, **b** HEK293T cells were co-transfected with V5-JAK1 or GST-V5 (**a**) and V5-TYK2 or GST-V5 (**b**) together with Flag-tagged PJA2-WT or PJA2-rm and HA-ubiquitin. Cells were lysed in a denaturing buffer containing 2 % SDS which was diluted to 0.7 % SDS prior to anti-V5 IP. Cell lysate and IP fractions were analyzed by SDS-PAGE and immunoblotting for the indicated proteins. Data are representative of at least four biologically independent experiments. **c** Quantification of the HA-ubiquitin signals in the experiments shown in panels **a** and **b**, with HA-ubiquitin quantities for the PJA2-WT condition made relative to the PJA2-rm condition. Bars represent means ± SDs from the four biologically independent replicates. P values were determined by one-way ANOVA and Dunnett’s T3 multiple comparisons. **d** Similar experiment to panel **b**, albeit an expression vector for untagged TYK2 was used. IPs were performed with anti-V5 or anti-TYK2 antibodies as indicated. Data are representative of three biologically independent experiments. **e**, **f** HEK293T cells were transfected with plasmids expressing Flag-mCherry or the indicated Flag-PJA2 variants (schematic in **f**) prior to generation of cell lysates and anti-Flag IP. Cell lysate and IP fractions were analyzed by SDS-PAGE and immunoblotting for the indicated proteins. Data are representative of three biologically independent experiments. PKA regulatory subunit binding domain (PKAR BD) and the catalytically active RING domain are indicated in **f**. **g**, **h** HEK293T cells were transfected with plasmids expressing V5-GFP or the indicated V5-TYK2 variants (schematic in **g**) prior to generation of cell lysates and anti-V5 IP. Cell lysate and IP fractions were analyzed by SDS-PAGE and immunoblotting for the indicated proteins. Data are representative of three biologically independent experiments. **i** HEK293T cells were co-transfected with V5-TYK2(1-588) together with Flag-tagged PJA2-WT or PJA2-rm and HA-ubiquitin. Cells were lysed in a denaturing buffer containing 2 % SDS which was diluted to 0.7 % SDS prior to anti-V5 IP. Cell lysate and IP fractions were analyzed by SDS-PAGE and immunoblotting for the indicated proteins. Data are representative of at least three biologically independent experiments. **j** HEK293T cells were co-transfected with V5-TYK2(1-588)-KR or GST-V5 together with Flag-tagged PJA2-WT or PJA2-rm and HA-ubiquitin. Cells were lysed in a denaturing buffer containing 2 % SDS which was diluted to 0.7 % SDS prior to anti-V5 IP. Cell lysate and IP fractions were analyzed by SDS-PAGE and immunoblotting for the indicated proteins. Data are representative of three biologically independent experiments. * indicates the IgG heavy or light chain visible on the immunoblot of the IP experiments.

TYK2 and JAK1 contain four domains: the N-terminal FERM and SH2 domains, which are both required for their interaction with IFNAR1/IFNAR2, and the C-terminal pseudo-kinase and kinase domains^72–74^. The TYK2 N-terminal FERM and SH2 domains together (residues 1-588) were sufficient to interact with PJA2 in truncation-based mapping studies (**Fig. 5g, h**). Furthermore, enhanced ubiquitination of this TYK2(1-588) construct, as well as the similar JAK1(1-582) construct, could occur in the presence of PJA2-WT (**Fig. 5i, Supplementary Fig. 4b**), suggesting that these domains are the predominant ubiquitination targets for PJA2. Canonical ubiquitination occurs on lysine residues^75,76^. To start mapping the ubiquitination site(s) on TYK2, we initially designed a TYK2(1-588) construct that completely lacks any lysine residues (including in the V5-tag) by substituting all 19 TYK2 lysines in this construct (as well as 1 lysine in the V5 tag) with arginine residues. Surprisingly, this lysine-less construct still exhibited enhanced ubiquitination in the presence of PJA2-WT (**Fig. 5j**), indicating that the ubiquitination site(s) on TYK2 are likely formed via a non-canonical ubiquitination of non-lysine residues, such as serine, threonine, and cysteine; or even to the N-terminal α-amino group of TYK2^77–80^. Overall, these data reveal that PJA2 can promote lysine-independent ubiquitination of TYK2, and likely JAK1, on its N-terminal FERM and SH2 domains.

### PJA2 promotes non-degradative ubiquitination of JAK1 and TYK2 and interferes with downstream STAT phosphorylation to negatively regulate type I IFN signaling

Ubiquitination is typically thought to lead to target protein degradation via the proteasome^75,76^. However, we found that siRNA-mediated depletion of PJA2 did not increase JAK1 or TYK2 protein levels following IFN-α2 stimulation (**Fig. 6a, Supplementary Fig. 5a, b**). Similarly, overexpression of PJA2 did not reduce JAK1 or TYK2 protein levels following IFN-α2 stimulation (**Fig. 6b**), nor lead to decreased levels of the Janus kinases when co-overexpressed (**Supplementary Fig. 5c**). We therefore reasoned that the non-lysine ubiquitination of TYK2 (and JAK1) promoted by PJA2 is probably via an atypical ubiquitin chain linkage (i.e. not via ubiquitin residue K48). This would be consistent with previous studies on PJA2 E3 ligase function, which have characterized it as forming non-degradative atypical polyubiquitin chains on some targets^71,81^. To understand mechanistically how PJA2 may negatively regulate IFN signaling without degrading TYK2/JAK1, we next assessed the impact of PJA2 on two well-described critical functions of these Janus kinases: regulation of IFN-α/β receptor levels^82^, and downstream STAT phosphorylation following IFN stimulation^83–85^. More precisely, TYK2 has previously been shown to stabilize IFNAR1 on the surface of unstimulated cells by preventing its internalization and turn-over via endocytosis^86^. Using CRISPR/Cas9, we generated several independent HEK293T-based TYK2 knockout (KO) cell lines and could confirm that lack of TYK2 reduces total levels of IFNAR1 (**Fig. 6c, d**). Furthermore, re-expression of TYK2 could partially restore IFNAR1 protein levels, but the further expression of PJA2-WT was unable to reverse this (**Fig. 6c, e**), suggesting that the ubiquitination of TYK2 promoted by PJA2 does not interfere with the ability of TYK2 to stabilize IFNAR1 levels. In contrast, we observed that siRNA-mediated depletion of PJA2 significantly increased the early IFN-α2-stimulated phosphorylation of STAT1 in both A549 and MRC-5 cells (**Fig. 6f-i**). Overall, these data are consistent with a working model whereby PJA2 interacts with the Janus kinases and acts as an E3 ubiquitin ligase to promote their non-lysine and non-degradative ubiquitination at N-terminal domains that functions to limit downstream signaling via the STATs to limit the response of cells to IFN (**Fig. 6j**).

**Fig. 6:**
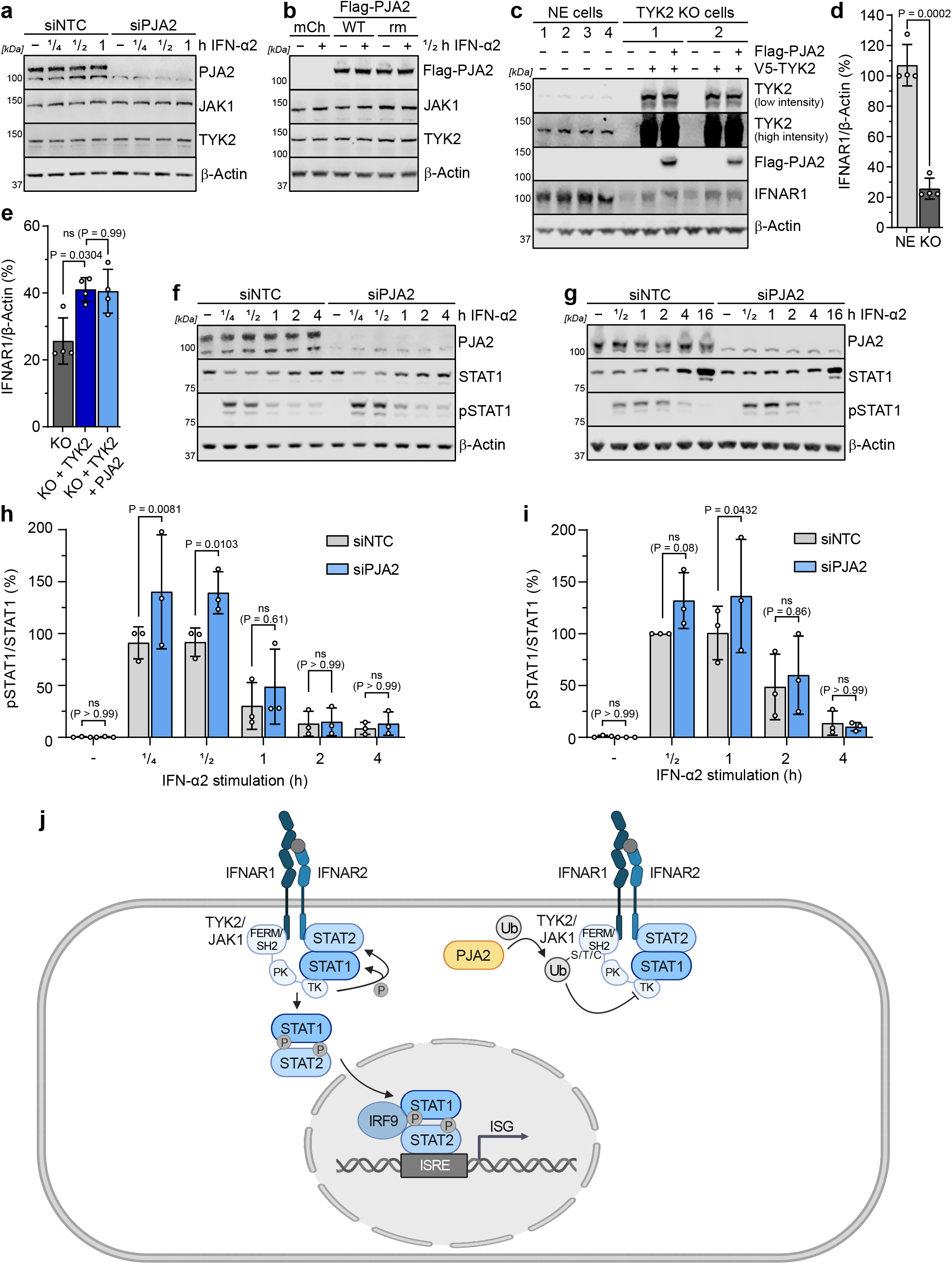
PJA2 promotes non-degradative ubiquitination of JAK1 and TYK2 and interferes with downstream STAT phosphorylation to negatively regulate type I IFN signaling. **a** MRC-5/hTERT cells were transfected with siRNA pools targeting PJA2 (or NTC) and were subsequently stimulated with 1000 IU/mL IFN-α2 for the indicated times. Total cell lysates were analyzed by SDS-PAGE and immunoblotting for the indicated proteins. Data are representative of three biologically independent experiments. **b** HEK293T cells were transfected with the indicated Flag-tagged construct and stimulated for 30 min with 1000 IU/mL IFN-α2. Total cell lysates were analyzed by SDS-PAGE and immunoblotting for the indicated proteins. Data are representative of at least two similar biologically independent experiments. **c** Non-edited HEK293T cell clones or TYK2-KO (knock out) cell clones were transfected with the indicated plasmids and total cell lysates were analyzed by SDS-PAGE and immunoblotting for the indicated proteins. Data are representative of at least two similar biologically independent experiments. **d**, **e** Quantification of IFNAR1 protein expression normalized to β-actin protein expression and made relative to one of the NE control cell clones in replicates of what is shown in panel **c**. Bars represent means ± SDs from two independent experiments with at least two independent NE and KO cell lines. Statistical significance was determined by one-way ANOVA and Dunnett’s T3 multiple comparisons. **f**, **g** A549 (**f**) or MRC-5/hTERT (**g**) cells were transfected with siRNA pools targeting PJA2 (or NTC) and were subsequently stimulated with 1000 IU/mL IFN-α2 for the indicated times. Total cell lysates were analyzed by SDS-PAGE and immunoblotting for the indicated proteins. Data are representative of three biologically independent experiments. **h**, **i** Quantification of pSTAT1 protein levels normalized to total STAT1 protein levels in replicates of what is shown in panels **f** (A549, **h**) and **g** (MRC-5/hTERT, **i**). Quantifications were made relative to a NTC siRNA transfected IFN-α2 stimulated sample in each experiment. Bars represent means ± SDs from three independent experiments. P values were determined by two-way ANOVA and Šídák’s multiple comparisons. Non-significant (ns) values are P > 0.05. **j** Working hypothesis integrating the findings from this study into a mechanistic model for how PJA2 can act as a negative regulator of IFN signaling.

## Discussion

Type I IFN signaling is an essential component of innate defenses against viral pathogens. This signaling cascade must be tightly regulated to prevent aberrant hyperactivation, yet still be able to react efficiently to limit viral disease. While key regulators of IFN signaling have previously been identified with different large-scale interaction or genetic approaches^5,33–39^, our current study describes the comprehensive system-wide application of TurboID-based proximity labeling and AP-MS to identify 103 proteins in close proximity to all seven core members of the human type I IFN signaling cascade: IFNAR1, IFNAR2, JAK1, TYK2, STAT1, STAT2, and IRF9. Importantly, analysis of this new proximal proteome unveiled an interconnected physical and functional network of factors and revealed previously unappreciated regulatory contributions to the human type I IFN pathway. Our approach is highly complementary to a similar BioID-based proximity labeling screen that was focused on identifying the proximal proteomes of murine STAT1, STAT2, and IRF9 in murine macrophage-like cells^51^. While species’ and cell-type differences likely explain the limited overlap in proximal proteins identified for STAT1, STAT2 and IRF9 in the two screens, it is striking that both screens did identify CNOT1 in proximity to the ISGF3 components. This factor is a member of the CCR4-NOT complex that negatively regulates type I IFN signaling^66,67^. Interestingly, two more components of the CCR4-NOT complex, CNOT2 and CNOT3, came up as repressors of ISG activation in a genetic screen^39^. As TurboID is a much more rapidly acting enzyme than BioID, allowing biotinylation within minutes rather than hours^43^, we were able to apply our system to study IFN-stimulated changes to the human signaling cascade at a very high temporal resolution, thereby providing new insights into the kinetics transient nature of the proximal factors we identified. Considering that the dataset we present here is the result of a comprehensive proximity labeling screen performed on all seven core members of the human type I IFN signaling cascade at the same time, and includes detailed information on interactions of known and unappreciated factors within this system, it should serve as a valuable future resource for characterizing novel effector mechanisms regulating this signaling cascade.

In our own follow up studies, we characterized the E3 ubiquitin ligase, PJA2, as a negative regulator of IFN signaling. PJA2 interacts with TYK2 and JAK1 and can promote their ubiquitination, strongly inhibiting IFN signaling by a mechanism that is not fully resolved, but which appears to at least include a partial limitation of downstream STAT1 phosphorylation. Ubiquitination of substrates by PJA2 often leads to their proteasomal degradation ^68–70,87^. However, there are also reports of PJA2 ubiquitinating proteins and impacting their functions without degradation. For example, PJA2 can ubiquitinate the HIV-1 Tat protein to stabilize Tat protein levels and to activate viral transcription^71^, and the PJA2-mediated ubiquitination of MFHAS1 can promote TLR2-mediated signaling^81^. Notably, we found that the ubiquitination of JAK1 and TYK2 promoted by PJA2 is independent of lysine residues in the Janus kinases and does not appear to lead to TYK2/JAK1 degradation. Recently, our knowledge on non-lysine ubiquitination events has greatly advanced and it is now a more broadly recognized mechanism to regulate protein functionality^88^. The non-lysine and non-degradative ubiquitination of TYK2/JAK1 could plausibly impact their conformation and/or interaction with other proteins. Indeed, given our observation that IFN-stimulated STAT1 phosphorylation increases following PJA2 depletion, it is likely that the ubiquitination of JAK1 and TYK2 at least partially impacts their conformation-dependent kinase activity. Janus kinases are known to switch between an autoinhibited (closed) and an open conformation in resting cells^89^. Following IFN binding to its receptor, the open conformation is required for JAK dimerization and activation^89^. We hypothesize that PJA2-mediated ubiquitination of JAK1 and TYK2 could stabilize their autoinhibited conformations or could act sterically to prevent (or limit) the kinases from dimerizing, thereby restraining their full activity.

We found PJA2 to be a constitutive interactor of TYK2 and JAK1, which opens up the question of whether, and how, the negative regulatory action of PJA2 itself may get activated. PJA2 does not appear to be an ISG^59^, unlike several other negative regulators of type I IFN signaling, such as USP18^59,90^ and SOCS1/3^59,91^. A previous study has shown that PJA2 can be activated by serine/threonine phosphorylation in its N-terminal region (S342/T389)^68^. It is possible that an unknown kinase can induce this phosphorylation as part of the IFN signaling response to activate a negative feedback loop. It is further possible that PJA2 acts constitutively to promote low-level ubiquitination of JAK1 and TYK2 to limit hyperactivity of the IFN system, and that this inhibitory ubiquitination must therefore be overcome to unleash the full signaling potential of the Janus kinases following IFN stimulation.

JAK1 and TYK2 are involved in JAK/STAT signaling pathways other than type I IFN signaling. Prominent examples are the other IFN signaling pathways, type II and type III IFN signaling, as well as IL-2, IL-6, and IL-10 cytokine signaling^92^. Here, we have already shown that PJA2 can affect not only type I, but also type II IFN signaling. Correspondingly, the ubiquitination of Janus kinases promoted by PJA2 might also play a broader role in several different signaling events. Interestingly, we identified one IL-6 family cytokine receptor complex, IL6ST/OSMR, proximal to both JAK1 and TYK2 in our TurboID screen, indicating that we might already have identified additional factors that are involved in other JAK/STAT signaling pathways. This may broaden the applicability of the dataset we provide here.

In summary, our comprehensive TurboID-based study has generated a robust dataset of > 100 proteins with high probability of being in close proximity to the core components of the human type I IFN signaling cascade, including IFNAR1, IFNAR2, JAK1, TYK2, STAT1, STAT2, and IRF9. This dynamic IFN-stimulated map of proteins in the proximity of these key signaling molecules led us to characterize an intriguing new mechanism by which non-lysine mediated ubiquitination may fine-tune the IFN pathway and its antiviral action. Our validated dataset should therefore form a solid basis for future investigations aimed at unraveling novel functional or regulatory intricacies of type I IFN signaling.

## Methods

### Cell culture

Human embryonic kidney (HEK) 293T cells and human lung epithelial A549 cells (both originally from ATCC) were cultured in Dulbecco’s Modified Eagle’s Medium (DMEM; Thermo Fisher Scientific), supplemented with 10 % fetal bovine serum (FBS) and 100 U/mL of penicillin-streptomycin (Gibco Life Technologies). hTERT-immortalized human lung fibroblasts (MRC-5/hTERT; a gift from Chris Boutell, University of Glasgow, UK) were cultured in Minimum Essential Medium Eagle (Sigma-Aldrich) supplemented with 10 % FBS, 100 U/mL of penicillin-streptomycin, 2 mM GlutaMAX (Thermo Fisher Scientific) and 1 % Non-Essential Amino Acids (NEAA; Thermo Fisher Scientific). Cells were maintained at 37 °C and 5 % CO_2_. IFN-α2 (NBP2-34971) and IFN-γ (NBP2-34992) were purchased from Novus Biologicals.

### Cloning of plasmids

The cDNA for TurboID^43^ was obtained from an existing vector (Addgene plasmid #107169; a gift from Alice Ting, Stanford University, USA) and cloned into pLVX-IRES-Puro (Takara) via *BamH*I and *BglI*I (V5-TurboID) or *Mfe*I and *EcoR*I (TurboID-V5). The V5-tag was added using appropriate primers. Full-length IFNAR1 (NM_174552.2, via *EcoR*I and *BamH*I), IFNAR2 (NM_000874.5, via *EcoR*I and *Not*I) and the Lyn11-plasma membrane localizing GFP control (via *EcoR*I and *BamH*I) were added to be N-terminal to the V5-TurboID coding sequence. JAK1 (NM_002227.4, via *Xho*I and *Not*I), TYK2 (NM_003331.5, via *EcoR*I and *Not*I), STAT1 (NM_007315.4, via *Not*I and *BamH*I), STAT2 (NM_005419.4, via *EcoR*I and *Not*I), IRF9 (NM_006084.5, via *EcoR*I and *BamH*I), and the other GFP controls (GFP, GFP-NES, GFP-NLS; via *EcoR*I and *BamH*I) were cloned to be C-terminal to the TurboID-V5 coding-sequence. cDNAs of STAT1, STAT2 and GFP were amplified from pre-existing vectors and the Lyn11-plasma membrane localization sequence^45^ (N-terminal), Rev NES^46^ (C-terminal) or SV40 large T antigen NLS^47^ (C-terminal) were cloned in-frame with the GFP coding sequence using appropriate primers. All other human IFN signaling component cDNAs were obtained by performing RT-PCR using total RNA from MRC-5/hTERT cells as a template. The HA-ubiquitin plasmid was obtained from Addgene (pRK5-HA-Ubiquitin-WT^93^, plasmid #17608, gift from Ted Dawson, Johns Hopkins University, USA). The p3xFlag-mCherry-CMV-7.1 and GST-V5 plasmids have been previously described^94,95^. PJA2 cDNA (NM_014819.5) was obtained from cellular mRNA by RT-PCR and cloned into p3xFlag-CMV-7.1 via *BamH*I/*BglI*I and *Not*I. V5-JAK1, V5-TYK2 (via *Not*I), and untagged TYK2 (via *EcoR*I and *Not*I) were cloned from the pLVX-TurboID-IRES-Puro vectors into pCDNA3.1. pCDNA3.1 containing the TYK2(1-588)-KR coding sequence was synthesized by GeneArt gene synthesis (Thermo Fisher Scientific). Specific nucleotide substitutions or coding-sequence truncations were introduced as required using the QuikChange XL Site-Directed Mutagenesis Kit (Agilent). All plasmid inserts and modifications were authenticated by Sanger sequencing.

### Generation of stable cell lines

To generate cell lines stably expressing TurboID-tagged proteins, lentiviral particles were first produced in HEK293T cells by co-transfection of the respective pLVX-TurboID-IRES-Puro construct, together with psPAX2 and pMD2.G (Addgene plasmids #12259 and #12260; gifts from Didier Trono, EPFL, Switzerland) at a ratio of 2:1:2. After 48h, lentivirus-containing supernatants were clarified by low speed centrifugation and filtration through a 0.45 μm filter prior to storage at −80°C. MRC-5/hTERT cells were subsequently transduced with the lentivirus-containing supernatants in the presence of polybrene (8 μg/mL, Sigma-Aldrich) and were selected with puromycin (2 μg/mL, Thermo Fisher Scientific) for at least 14 days.

### Proximity labeling

MRC-5/hTERT cells expressing the appropriate TurboID-tagged protein were split 1:4 into 10 cm dishes and left to grow for 5 days until confluent. Cells were then serum-starved for 20 h before treatment with 1000 IU/mL IFN-α2 for 0 min, 20 min, 1 h or 2 h. During the last 15 min, biotin (Sigma-Aldrich) was added to a final concentration of 500 μM. Cells were subsequently washed five times with PBS at 4 °C and lysed in RIPA buffer (50 mM Tris-HCl pH 8, 150 mM NaCl, 0.1 % SDS, 0.5 % sodium deoxycholate, 1 % Triton X-100) supplemented with cOmplete Mini EDTA-free protease inhibitors (Roche). Lysates were sonicated, cleared by centrifugation at 10,000 x g for 15 min, and a fraction was removed for analysis of the total cell lysate. After incubation of the remaining lysate with streptavidin magnetic beads (Thermo Fisher Scientific) for 16 h at 4 °C with end-over-end tumbling, samples were washed twice with RIPA buffer, once with 1 M KCl, once with 0.1 M Na_2_CO_3_, once with 1 M urea in 10 mM Tris-HCl (pH 8.0), and twice in freshly prepared 50 mM ammonium bicarbonate. Mass spectrometry was performed by the Functional Genomics Center Zurich and data were analyzed using SAINTexpress as previously described^5,49^. Specific selection criteria were a fold change (FC) ≥ 4 in protein abundance (label-free quantification) over the similarly localized GFP control, as well as a FC ≥ 2 over all other GFP controls, and an interaction probability (SAINT probability) ≥ 0.7 with at least 2 unique spectral counts. The mass spectrometry proteomics data have been deposited to the ProteomeXchange Consortium via the PRIDE^96^ partner repository with the dataset identifier PXD045482. The data will be released following peer-review.

### Transfections and RNA interference

For plasmid transfections, 70 % confluent HEK293T cells were transfected using FuGENE HD transfection reagent (Promega) according to the manufacturer’s protocol. When siRNAs and plasmids were co-transfected into HEK293T cells, reverse transfection with Lipofectamine 2000 (Thermo Fisher Scientific) was performed according to the manufacturer’s protocol. For siRNA-mediated knockdown experiments, MRC-5/hTERT or A549 cells were reverse transfected with ON-TARGETplus siRNA SMARTpools (Horizon Discovery) or FlexiTube siRNA (Qiagen) using Lipofectamine RNAiMAX transfection reagent (Thermo Fisher Scientific) according to the manufacturer’s protocol. Detailed information on the siRNAs used in this study can be found in **Supplementary Tables 1 and 2**.

### VSV-GFP replication assay

MRC-5/hTERT or A549 cells were reverse transfected with siRNAs in 96-well plates for 32 h or 44 h, before stimulation with 250 IU/mL IFN-α2 for a further 16 h or 4 h. Cells were then infected with VSV-GFP (MOI = 1 PFU/cell; gift from Peter Palese, Icahn School of Medicine, New York, USA) in FluoroBrite DMEM (Thermo Fisher Scientific) supplemented with 5 % FBS, and cells were monitored with the IncuCyte live-cell imaging system (Sartorius) for up to 4 days. Total green object integrated intensities (Green Calibrated Unit (GCU) x µm²/Image) were exported and the area under the curve was calculated in GraphPad Prism 9.

### RT-qPCR analysis

RNA was isolated with the ReliaPrep RNA Cell Miniprep kit (Promega) according to the manufacturer’s protocol. 250 ng RNA were reverse transcribed into cDNA with SuperScript III (Thermo Fisher Scientific) and an Oligo(dT)_15_ primer (Promega). qPCR was performed with the Fast EvaGreen qPCR Master Mix (Biotium) and measured on a ABI7300 Real-Time PCR system (Applied Biosystems) in duplicates in 96-well plates. Specific cDNA quantities were calculated based on the ΔΔC_t_ method and normalized to GAPDH as a housekeeping gene. The primers used are listed in **Supplementary Table 3**.

### Immunoprecipitation (IP)

For standard co-IP experiments, cells were washed once with PBS and lysed in co-IP lysis buffer (50 mM Tris HCl pH 7.5, 150 mM NaCl, 1 mM EDTA, 1 % Triton X-100) supplemented with cOmplete Mini EDTA-free protease inhibitors (Roche). Lysates were then sonicated, and cleared by centrifugation at 16,000 x g for 15 min. For denaturing IP experiments, cells were washed once with PBS and lysed in a denaturing SDS lysis buffer (50 mM Tris-HCl pH 7.8, 650 mM NaCl, 1 % NP-40, 2 % SDS, 5 mM EDTA, 20 mM N-ethylmaleimide, 10 mM β-mercaptoethanol) supplemented with cOmplete Mini EDTA-free protease inhibitors (Roche). After sonication, lysates were diluted 1:3 in SDS dilution buffer (50 mM Tris-HCl pH 7.8, 650 mM NaCl, 1 % NP-40, 0.1 % SDS, 5 mM EDTA, 20 mM N-ethylmaleimide) supplemented with cOmplete Mini EDTA-free protease inhibitors (Roche), and centrifuged at 20,000 x g for 30 min. For all IPs, a sample of the cleared lysate was removed for analysis of the total cell lysate. Cleared lysates were then incubated with the indicated antibodies at 4 °C for 16 h with end-over-end tumbling. Protein G Dynabeads (Thermo Fisher Scientific) were added for 30 min at room temperature with further tumbling. Standard co-IP samples were subsequently washed three times with co-IP lysis buffer, while denaturing IP samples were washed six times with SDS dilution buffer. Immunoprecipitated proteins and total cell lysates were analyzed by immunoblotting.

### Immunoblotting

Samples were diluted in 4x Laemmli protein sample buffer (Bio-Rad) containing 10 % β-mercaptoethanol, or directly lysed in 1x Laemmli buffer, and sonicated to shear DNA. Proteins were separated on Bolt 4 – 12 % Bis-Tris Plus Gels (Thermo Fisher Scientific) and transferred onto 0.45 μm nitrocellulose membranes according to the manufacturer’s protocol. Membranes were then blocked with 5 % milk or 5 % BSA in TBS-T, and incubated with the appropriate primary antibody: β-actin (Santa Cruz, sc-47778), actin (Sigma-Aldrich, A2103), Flag (Sigma-Aldrich, F1804 or F7425), V5 (Bio-Rad, MCA1360; or Bethyl Laboratories, A190-119A), GST-HRP conjugate (Cyriva, RPN1236), HA (Cell Signaling Technology, 2367 or 3724), IFNAR1 (Abcam, ab124764), JAK1 (BD Biosciences, 610231), TYK2 (Cell Signaling Technology, 14193), PJA2 (Bethyl Laboratories, A302-991A), STAT1 (Santa Cruz, sc-417), phospho-Y701-STAT1 (Cell Signaling Technology, 9167), STAT2 (Santa Cruz, sc-1668), phospho-Y690-STAT2 (Cell Signaling Technology, 88410), MxA (ab143^97^; gift from Jovan Pavlovic, University of Zurich, Switzerland), IRF9 (BD Biosciences, 610285), DNAJA2 (Abcam, ab157216), KLC2 (Atlas Antibodies, HPA040416), CNOT1 (Cell Signaling Technology, 44613), or IL6ST (Bethyl Laboratories, A304-929A-T). After subsequent washing, membranes were incubated with the appropriate secondary antibody: IRDye 800CW Goat anti-Mouse IgG (LI-COR Biosciences, 926-32210), IRDye 800CW Goat anti-Rabbit IgG (LI-COR Biosciences, 926-32211), IRDye 800CW Donkey anti-Goat IgG (LI-COR Biosciences, 925-32214), IRDye 680RD Goat anti-Mouse IgG (LI-COR Biosciences, 926-68070), IRDye 680RD Goat anti-Rabbit IgG (LI-COR Biosciences, 926-68071), HRP-Horse anti-Mouse IgG (Cell Signaling Technology, 7076P2), HRP-Goat anti-Rabbit IgG (Sigma-Aldrich, A0545), or HRP-Rabbit anti-Goat IgG (Sigma-Aldrich, A4174). The signal was detected with the Odyssey Fc Imager (LI-COR Biosciences) and quantified with the Image Studio Lite Quantification software (LI-COR Biosciences).

### Immunofluorescence microscopy

Cells were seeded on coverslips in 24-well plates for 24 h and, where indicated, were stimulated with 1000 IU/mL IFN-α2. Cells on coverslips were then washed once with PBS, fixed with 3.7 % paraformaldehyde for 10 min at room temperature, washed three more times with PBS, and then permeabilized with 100 % methanol for 10 min at −20 °C. After three more washes in PBS, samples were blocked in 2 % FBS in PBS and incubated with the indicated primary antibodies (see immunoblotting section) in 2 % FBS in PBS. DNA was stained with DAPI (Sigma-Aldrich). Secondary antibodies were incubated with the samples in PBS supplemented with 2 % FBS. The secondary antibodies used were Alexa Fluor 488/555 anti-mouse/rabbit (Thermo Fisher Scientific, A11029, A21206, A31570 or A31572). Coverslips were mounted with ProLong Gold Antifade (Thermo Fisher Scientific) and imaged with a Leica DM IL LED microscope (Leica Microsystems) using the LasX software.

### Luciferase reporter assay

In a 96-well plate, HEK293T cells were reverse transfected with a Firefly luciferase-based reporter plasmid (see below), a constitutively active Renilla luciferase control (pRL-TK-Renilla) and various amounts of pCMV7.1-3xFlag-mCherry, -PJA2-WT, or -PJA2-rm. Total plasmid levels were kept constant by addition of an appropriate amount of empty pCMV7.1-3xFlag vector. Firefly luciferase was expressed either under control of the murine *Mx1* promoter (pGL3-Mx1P-FFluc^98^; gift from Georg Kochs, University of Freiburg, Germany), the human *ISG54* promoter (pISG54-Luc^99^; gift from Chris Basler, Icahn School of Medicine, New York, USA), or 3 copies of the IFN-γ activation site (GAS; pGAS-Luc^99^; gift from Chris Basler, Icahn School of Medicine, New York, USA). 30 h post-transfection, cells were stimulated (or mock) with 100 or 1000 IU/mL of IFN-α2 or 1000 IU/mL of IFN-γ for 16 h. Cells were then lysed and Firefly and Renilla luciferase activities were determined with the Dual-Luciferase Reporter Assay System (Promega) and an EnVision plate reader (PerkinElmer) according to the manufacturers’ protocols.

### Generation of knockout cells

HEK293T knockout cells were generated using a ribonucleoprotein (RNP) system as described previously^100^. Briefly, HEK293T cells were reverse transfected with preassembled RNP complexes using Lipofectamine RNAiMAX transfection reagent (Thermo Fisher Scientific). Knockout cell clones were obtained by limiting dilution and verified via NGS and immunoblotting. crRNA and NGS primer sequences are listed in **Supplementary Tables 4 and 5**.

### Gene ontology analyses

Functional enrichment gene ontology terms^56,57^ were analyzed with DAVID^63^ using all human genes as background. Bait proteins were excluded from the analyses.

### Network generation

Protein landscape networks were generated using Cytoscape 3.9^101^. Proteins belonging to the viral process (GO:0016032)^56,57^ category (**Fig. 2**) were annotated with Cytoscape 3.9^101^.

### Statistical analyses

Statistical analyses were performed in GraphPad Prism 9. Data were analyzed by two-way ANOVA and Tukey’s multiple comparisons (luciferase reporter assay) or Šídák’s multiple comparisons (immunoblot quantification of siRNA knockdowns, qPCR quantifications, VSV-GFP assay), or one-way ANOVA and Dunnett’s T3 multiple comparisons (immunoblot quantification of HA-ubiquitin ratios, IFNAR1 expression levels).

## Acknowledgements

We are grateful to Davide Eletto (ETHZ, Switzerland) for providing important technical insights at the instigation of this project. We are also grateful to Jovan Pavlovic (University of Zurich, Switzerland), Chris Boutell (University of Glasgow, UK), Alice Ting (Stanford University, USA), Ted Dawson (Johns Hopkins University, USA), Didier Trono (EPFL, Switzerland), Peter Palese (Icahn School of Medicine, New York, USA), Chris Basler (Icahn School of Medicine, New York, USA) and Georg Kochs (University of Freiburg, Germany) for kind gifts of antibodies, cells, plasmids and viruses. Mass spectrometry experiments were performed at the Functional Genomics Center Zurich (FGCZ) of the University of Zurich and the ETH Zurich. Imaging was performed with equipment maintained by the Center for Microscopy and Image Analysis, University of Zurich, Switzerland. Schematic figures were created with BioRender.com. The research leading to these results received funding from the Swiss National Science Foundation (grants 31003A_182464 and 310030_214957 to BGH).

## Author contributions

Conceptualization: SS and BGH; Methodology, Validation, Formal Analysis, and Investigation: SS; Visualization: SS; Writing – Original Draft: SS; Writing – Review & Editing: BGH; Supervision: BGH; Funding Acquisition and Project Administration: BGH.

