## Supplemental Tables and Figures for "Proximal protein landscapes of the type I interferon signaling cascade reveal negative regulation by the E3 ubiquitin ligase PJA2"

### Supplementary information

#### Supplementary Table 1: Qiagen FlexiTube siRNAs

| Target | Catalog Number | Gene Accession |
|---|---|---|
| **AllStars negative control** | 1027280 | |
| **IFNAR1** | SI00013195 | NM_000629 |
| **IFNAR2** | SI05005896 | NM_207585 |
| **JAK1** | SI04991378 | NM_002227 |
| **TYK2** | SI02223221 | NM_003331 |
| **STAT1** | SI05078556 | NM_007315, NM_139266 |
| **STAT2** | SI05020876 | NM_005419, NM_198332 |
| **IRF9** | SI00084364 | NM_006084 |

#### Supplementary Table 2: Horizon Discovery ON-TARGETplus siRNA SMARTpools

| Target | Catalog Number | Gene Accession |
|---|---|---|
| **ON-TARGETplus Non-targeting Control** | D-001810-10 | |
| **AFF1** | L-020074-02 | NM_001166693 |
| **AIM1** | L-024709-01 | NM_001624 |
| **ALDH1B1** | L-008254-00 | NM_000692 |
| **ALMS1** | L-012889-00 | NM_015120 |
| **ANKRD17** | L-013554-01 | NM_198889 |
| **APC** | L-003869-00 | NM_000038 |
| **APPL1** | L-005138-00 | NM_012096 |
| **ARFGEF1** | L-012207-00 | NM_006421 |
| **ASCC2** | L-016458-02 | NM_032204 |
| **CALCOCO1** | L-007038-01 | NM_020898 |
| **CNOT1** | L-015369-01 | NM_206999 |
| **CTPS1** | L-006644-00 | NM_001905 |
| **DNAJA2** | L-012104-00 | NM_005880 |
| **DNMBP** | L-026304-01 | NM_015221 |
| **ECD** | L-019678-00 | NM_007265 |
| **EEF1D** | L-011648-01 | NM_001130056 |
| **EIF4E2** | L-019870-01 | NM_004846 |
| **FAM175B** | L-016146-01 | NM_032182 |
| **GIGYF2** | L-013918-01 | NM_015575 |
| **HECTD1** | L-007188-00 | NM_015382 |
| **HSP90AB1** | L-005187-00 | NM_007355 |
| **HTRA1** | L-006009-00 | NM_002775 |
| **IFNAR2** | L-015411-00 | NM_207584 |
| **IGF2BP3** | L-003976-00 | NM_006547 |

| | | |
|---|---|---|
| **IL6ST** | L-005166-00 | NM_175767 |
| **IRF9** | L-020858-00 | NM_006084 |
| **KLC1** | L-019482-00 | NM_182923 |
| **KLC2** | L-014218-00 | NM_022822 |
| **LRBA** | L-012751-00 | NM_006726 |
| **LRCH4** | L-011321-01 | NM_002319 |
| **MADD** | L-004429-00 | NM_130474 |
| **MAP1A** | L-013482-00 | NM_002373 |
| **MFSD10** | L-016015-01 | NM_001120 |
| **MTCL1** | L-023376-01 | NM_015210 |
| **MYCBP2** | L-006951-00 | NM_015057 |
| **NT5E** | L-008217-00 | NM_002526 |
| **OSMR** | L-008050-00 | NM_003999 |
| **PCCA** | L-008965-00 | NM_000282 |
| **PCM1** | L-005165-00 | NM_006197 |
| **PGM3** | L-013912-01 | NM_015599 |
| **PHLDB2** | L-016702-01 | NM_145753 |
| **PJA2** | L-006916-00 | NM_014819 |
| **PLEKHG2** | L-023690-00 | NM_022835 |
| **RAB1B** | L-008958-01 | NM_030981 |
| **RIPK1** | L-004445-00 | NM_003804 |
| **SOCS3** | L-004299-00 | NM_003955 |
| **STAT6** | L-006690-00 | NM_003153 |
| **TBC1D2** | L-020463-01 | NM_018421 |
| **TEX2** | L-017117-02 | NM_018469 |
| **TMEM131** | L-022532-02 | NM_015348 |
| **USP18** | L-004236-00 | NM_017414 |
| **USP9X** | L-006099-00 | NM_021906 |
| **WNK1** | L-005362-02 | NM_014823 |
| **WWC2** | L-016585-02 | NM_024949 |

**Supplementary Table 3: qPCR primers**

| Target | Forward primer | Reverse primer |
|---|---|---|
| **ISG54** | GCGTGAAGAAGGTGAAGAGG | GCAGGTAGGCATTGTTTG |
| **MX1** | AGACAAGGTTGTGGACGTGG | TTCCTCCAGCAGATCCCTGA |
| **GAPDH** | CTGGCGTCTTCACCACCATGG | CATCACGCCACAGTTTCCCGG |

**Supplementary Table 4: crRNA**

| Target | Sequence |
|---|---|
| **TYK2** | ACCUGCGGAAGACGUUCCGAGUUUUAGAGCUAUGCU |

**Supplementary Table 5: NGS primers**

| Target | Forward primer | Reverse primer |
|---|---|---|
| **TYK2** | CTTTCCCTACACGACGCTCTTCCGA<br>TCTCCCAGCTTCAAGGACTGCAT | GACTGGAGTTCAGACGTGTGCTCTTC<br>CGATCTCACTGTCCCGGATGTAGCAG |

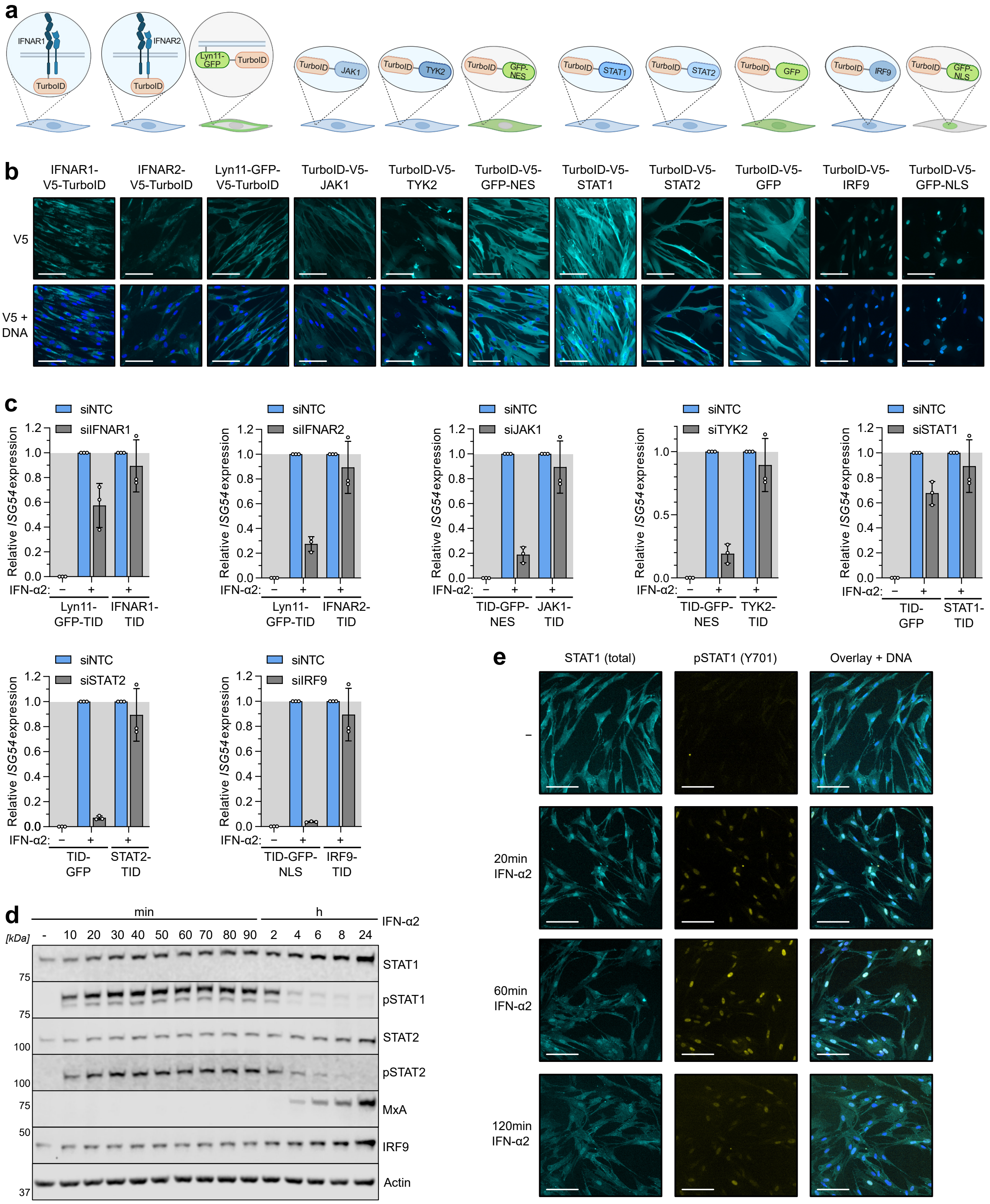


**Supplementary Fig. 1: Validation of TurboID-based type I IFN signaling constructs.**

**a** Schematic representation of the TurboID-tagged constructs used for proximity labeling. Each construct has a V5 tag (not depicted) that serves as a linker between TurboID and the protein of interest. **b** Immunofluorescence (IF) imaging of the intracellular localization of each TurboID-tagged construct in the transduced MRC-5/hTERT cells using anti-V5 antibody. DNA was stained with DAPI. Scale bars represent 100 µm.

**c** *ISG54* mRNA expression as determined by RT-qPCR following ± 4 h IFN-α2 stimulation (1000 IU/mL). The indicated transduced MRC-5/hTERT cells had previously been reverse transfected with siRNAs targeting the 3'untranslated region of the indicated gene or a non-targeting siRNA (NTC). Data are normalized to GAPDH expression levels in the same sample and made relative to the NTC + IFN-α2 condition. Bars represent means ± SDs from three biologically independent experiments conducted in technical duplicates.

**d** MRC-5/hTERT cells were stimulated with 1000 IU/mL IFN-α2 for the indicated times. Total cell lysates were analyzed by SDS-PAGE and immunoblotting for the indicated proteins.

**e** IF imaging of total and phosphorylated STAT1 in MRC-5/hTERT cells following stimulation with 1000 IU/mL IFN-α2 for the indicated times. DNA was stained with DAPI. Scale bars represent 100 μm.

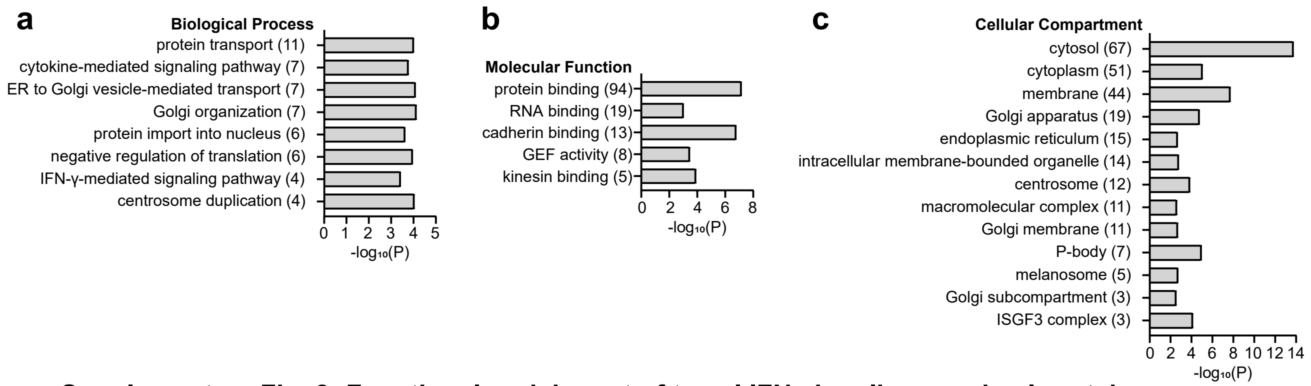


**Supplementary Fig. 2: Functional enrichment of type I IFN signaling proximal proteins.**
Functional enrichment of biological process (**a**), molecular function (**b**), or cellular compartment (**c**) gene ontology terms of the identified type I IFN signaling proximal proteins analyzed with DAVID. $-\log_{10}$(P value) is depicted for all false discovery rate < 0.05 terms. The number in brackets indicates the number of hits per category.

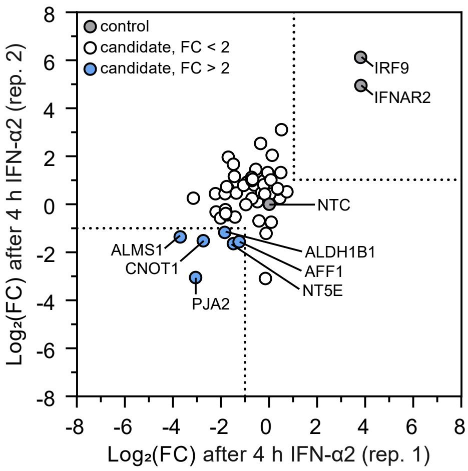


**Supplementary Fig. 3: siRNA screening identifies proximal proteins that functionally regulate type I IFN antiviral activity.**

$Log_2$(FC) in VSV-GFP replication (AUC values) after 4 h IFN-α2 stimulation of cells previously transfected with siRNAs targeting the indicated genes. Log2(FC) is relative to VSV-GFP replication in the NTC condition. Two biologically independent replicates are plotted. The dotted lines indicate a 2-fold change from the VSV-GFP replication in the NTC condition.

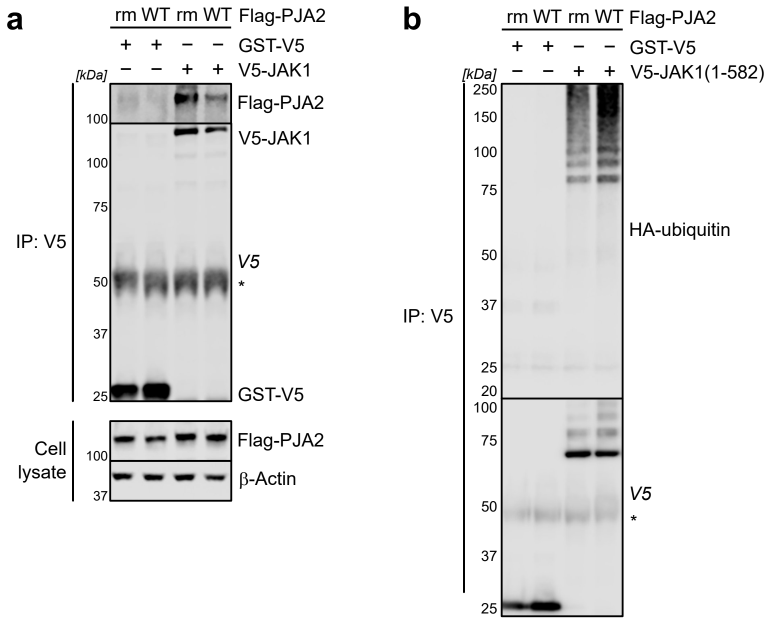


**Supplementary Fig. 4: PJA2 interacts with JAK1 and promotes the ubiquitination of JAK1(1-582).**
**a** HEK293T cells were transfected with plasmids expressing V5-JAK1(1-582) or GST-V5 together with Flag-PJA2-WT or PJA2-rm to generate cell lysates prior to anti-V5 immunoprecipitation (IP). Cell lysate and IP fractions were then analyzed by SDS-PAGE and immunoblotting for the indicated proteins. Data are representative of at least two biologically independent experiments.
**b** HEK293T cells were co-transfected with V5-JAK1(1-582) or GST-V5 together with Flag-tagged PJA2-WT or PJA2-rm and HA-ubiquitin. Cells were lysed in a denaturing buffer containing 2 % SDS which was diluted to 0.7 % SDS prior to anti-V5 IP. Cell lysate and IP fractions were analyzed by SDS-PAGE and immunoblotting for the indicated proteins. Data are representative of at least two biologically independent experiments. * indicates IgG heavy chain.

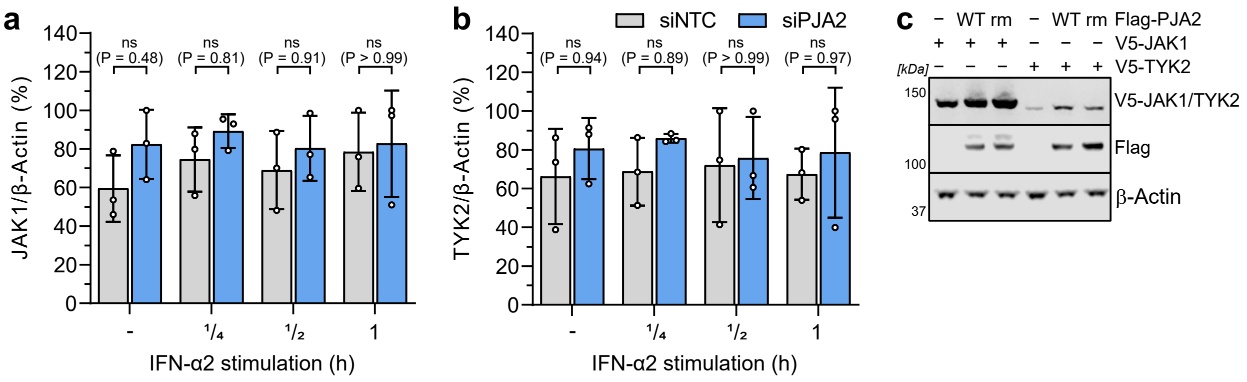


**Supplementary Fig. 5: PJA2 does not affect JAK1 and TYK2 expression levels.**

**a**, **b** Immunoblot quantification of JAK1 (**a**) or TYK2 (**b**) protein levels normalized to β-Actin protein levels in replicate experiments of that shown in main Fig. 6a. Quantifications were made relative to the highest value in each experiment. Bars represent means ± SDs from three independent experiments. P values were determined by two-way ANOVA and Šídák's multiple comparisons. Non-significant (ns) values are $P > 0.05$.

**c** HEK293T cells were co-transfected with V5-JAK1 or V5-TYK2 together with Flag-tagged PJA2-WT, Flag-tagged PJA2-rm or empty vector control. Total cell lysates were analyzed by SDS-PAGE and immunoblotting for the indicated proteins.
